# Parabrachial *Calca* neurons drive nociplasticity

**DOI:** 10.1101/2023.10.26.564223

**Authors:** Logan F. Condon, Ying Yu, Sekun Park, Feng Cao, Jordan L. Pauli, Tyler S. Nelson, Richard D. Palmiter

## Abstract

Pain that persists beyond the time required for tissue healing and pain that arises in the absence of tissue injury are poorly understood phenomena mediated by plasticity within the central nervous system. The parabrachial nucleus (PBN) is a hub that relays aversive sensory information and appears to play a role in nociplasticity. Here, by preventing PBN *Calca* neurons from releasing neurotransmitter or directly stimulating them we demonstrate that activation of *Calca* neurons is both necessary for the manifestation of chronic pain after nerve ligation and is sufficient to drive nociplasticity in wild-type mice. Aversive stimuli such as exposure to nitroglycerin, cisplatin, or LiCl can drive nociplasticity in a *Calca*-neuron-dependent manner. Calcium fluorescence imaging reveals that nitroglycerin activates PBN *Calca* neurons and potentiates their responses to mechanical stimulation. The activity and excitability of *Calca* neurons increased for several days after aversive events, but prolonged nociplasticity likely occurs in downstream circuitry.

## INTRODUCTION

Pain is an evolutionarily adaptive sensory modality that signals tissue injury and triggers defensive behavioral responses. However, changes in the neural circuitry underlying pain sensation can drive pain in the absence of tissue injury, a debilitating condition referred to as nociplastic pain (Fitzcharles et al., 2021; Nijs et al 2021). While nociplastic pain can occur independently, it often presents as part of a mixed-pain pathophysiology, arising in parallel with chronic nociceptive or neuropathic pain (Fitzcharles et al., 2021; Kosek et al., 2021; Nijs et al 2021; Maixner et al., 2016). This suggests that persistent pain experience can drive changes in pain perception and produce a generalized pain state. Chronic pain affects up to one in five people (Mills et al., 2018; Dahlhamer et al., 2018; Maixner et al., 2016) and a nociplastic component is estimated to be present in 25% to 75% of chronic pain cases (Dydyk et al., 2023; Fitzcharles et al., 2021). The neural circuitry and neuroplasticity underlying nociplastic pain are not well understood. Interrogating the pathophysiology of this widespread condition is necessary to develop therapeutic interventions that target the mechanistic underpinnings of nociplastic pain, not just the presenting symptoms.

Nociceptive signaling from the spinal cord, trigeminal and vagal nerves, and area postrema can activate PBN neurons (Choi et al., 2021; Rodriguez et al., 2017; Hermann et al., 1985; Zhang et al., 2021), which relay those signals to multiple forebrain regions (Pauli et al., 2022; Huang et al., 2021; Chiang et al., 2020; Grady et al., 2020; Saper et al., 1980; Gauriau and Bernard, 2002). The PBN is composed of approximately a dozen molecularly defined glutamatergic neuron subtypes and a minor population of GABAergic neurons (Pauli et al. 2022). Non-specific activation of the PBN glutamatergic neurons can elicit painful phenotypes such as allodynia, while inhibition of those neurons or activation of the GABAergic neurons can inhibit painful phenotypes (Torres-Rodriguez et al., 2023; Zhou et al., 2023; Sun et al., 2020; Chiang et al., 2019). The roles that molecularly defined subsets of PBN neurons contribute to pain sensation is also being explored. For example, chemogenetic activation of PBN *Tac1*-expressing neurons elicits escape-like behaviors (Arthurs, et al. 2023; Barik et al., 2018), whereas chemogenetic or optogenetic activation of *Tacr1*-expressing neurons elicits coping responses to painful stimuli (Barik et al., 2021; Huang et al., 2021; Ma 2021; Deng, et. al., 2020). We and others have shown that PBN *Calca* neurons are activated by aversive stimuli spanning several modalities including some that are considered painful, e.g., foot shock, tail pinch, and formalin- or Freund’s adjuvant-induced inflammation (Kang et al., 2022; Campos et al., 2018; Chen et al., 2018; Campos et al. 2017; Han et al., 2015).

Chemogenetic activation of *Calca* neurons promotes anorexia, adipsia, and escape-like behaviors (Arthurs et al., 2023; Carter et al., 2013) whereas more robust stimulation with optogenetics can induce freezing, bradycardia, and fear-like behaviors (Bowen et al., 2020; Han et al., 2015). Most assays have examined effects of transient activation, but chronic activation can promote severe anorexia (Carter et al., 2013) and the neurological effects of cancer cachexia have been shown to depend on activation of PBN *Calca* neurons (Campos et al., 2017). However, the effects of chronic activation of *Calca* neurons on pain-related phenotypes have not been examined. We show here that activation of *Calca* neurons is necessary to establish allodynia after nerve ligation. Additionally, chronic activation of these neurons is sufficient to instate chronic pain phenotypes, suggesting that the activity of *Calca* neurons drives nociplasticity.

## RESULTS

### Parabrachial *Calca* neurons are necessary for the induction of neuropathic pain

Chronic pain was modeled using unilateral, partial sciatic nerve ligation (pSNL) (Malmberg and Basbaum, 1998) and the presence of neuropathic allodynia was confirmed using the von Frey tactile-sensitivity assay. pSNL produced a significant reduction in the paw-withdrawal threshold both ipsilateral and contralateral to the site of nerve injury (Fig. S1A), consistent with previous findings of bilateral tactile hypersensitivity after nerve injury (Abraham et al., 2020; Raver et al., 2020; Arguis et al., 2008; Koltzenburg et al., 1999). PBN *Calca* neurons are known to act as a primary relay for brief pain signals (Campos et al., 2018; Palmiter, 2018; Han et al., 2015); however, it was not known whether they are involved in the experience of chronic pain.To determine whether they are activated in chronic pain we performed RNAscope *in situ* hybridization on sham or pSNL-operated mice using probes for *Calca, Cck* and the immediate early gene *Fos*, a proxy for neuronal activity (Chung, 2015) (Fig, 1A). We included *Cck* because it was shown to be reduced 30 days following pSNL (Fu et al., 2022) and could serve as a positive control. Three days following pSNL, we observed a significant increase in the colocalization of *Fos* and *Calca* mRNA in the PBN of pSNL-treated animals; however, this effect did not persist at 30 days post-surgery (dps) (Fig. 1B, F). There was no difference in the colocalization of *Calca* and *Fos* at either time point when comparing the ipsilateral and contralateral PBN, or when comparing males and females (Fig. S1B, C). Additionally, there was a significant increase in the *Fos* induction of Calca-negative PBN neurons at the 3 dps, but not the 30 dps (Fig. 1C, G). There was no change in the expression level of *Calca* within the PBN when comparing sham and pSNL-treated animals at either time point (Fig. 1D, H). However, the percentage of *Cck*-positive neurons was significantly decreased at 30 dps (Fig. 1E, I), consistent with a previous finding (Fu et al., 2022). These data suggest that *Calca* neurons are activated by pSNL, but the activity subsequently wanes suggesting that it is not involved in the maintenance of chronic pain.

**Figure 1.**
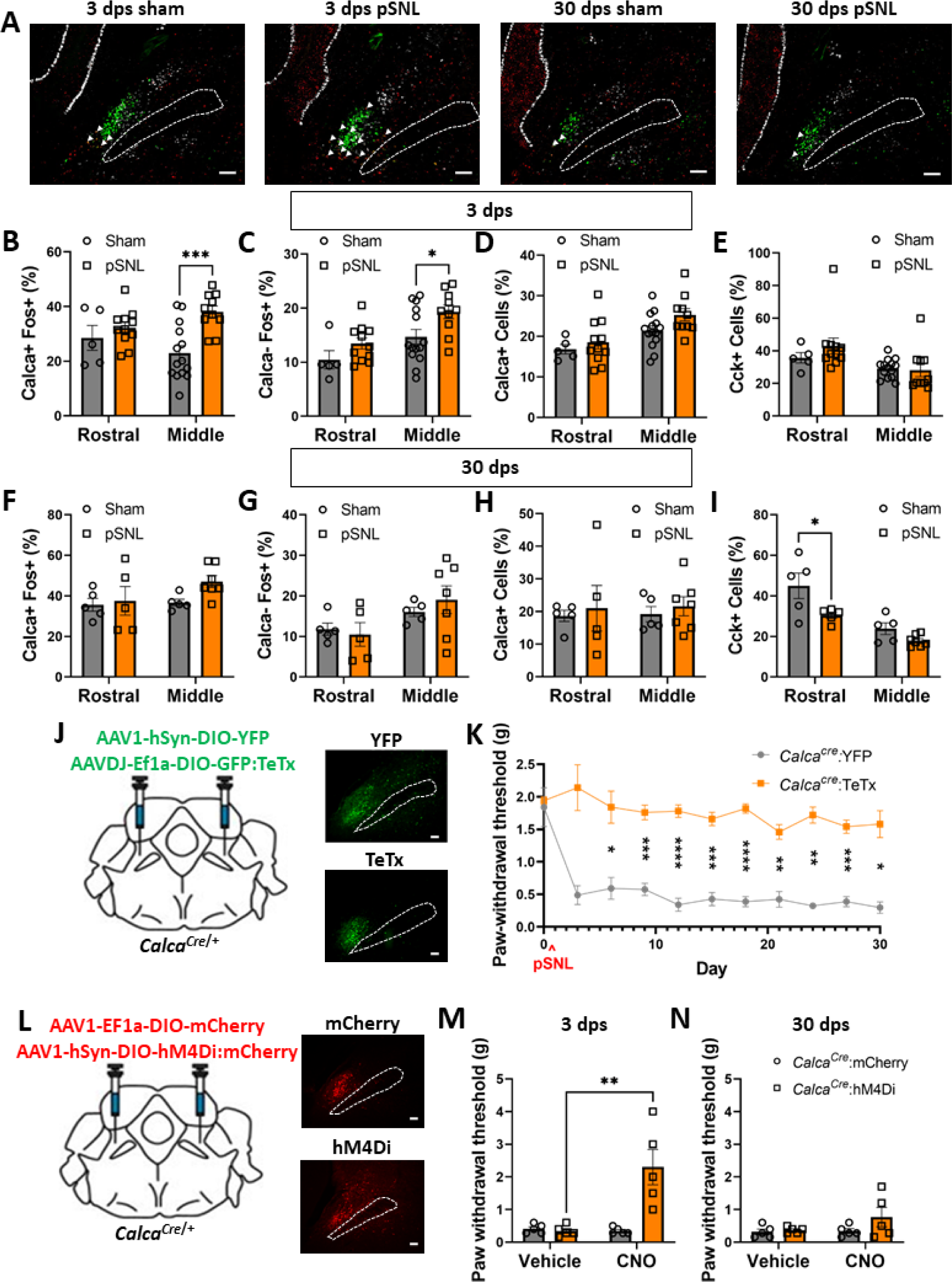
Parabrachial *Calca* neurons are necessary for the induction of neuropathic pain. (A) Representative images from RNAscope *in situ* hybridization on tissue slices collected 3 or 30 days post sham or pSNL surgery with probes targeting *Calca* (green), *Fos* (red), and *Cck* (white). Scale bar = 100 µm, dotted line marks the superior cerebellar peduncle (SCP), anterior-posterior Bregma level = -5.1. (B) pSNL increased *Fos* mRNA in *Calca* neurons in the middle, but not the rostral, PBN at 3 dps. (C) pSNL increased *Fos* mRNA in non-*Calca* neurons in the middle, but not the rostral, PBN at 3 dps. (D) pSNL did not change the number of *Calca*-positive neurons in the rostral or middle PBN at 3 dps. (E) pSNL did not change the number of *Cck* positive cells in the rostral or middle PBN at 3 dps. (B-E) Rostral sham n = 5, middle sham n = 14, rostral pSNL n = 11, and middle pSNL n = 10. (F) pSNL did not drive an increase in the expression of *Fos* mRNA in *Calca* neurons at 30 dps. (G) pSNL did not drive an increase in the expression of *Fos* mRNA in non-*Calca* neurons at 30 dps. (H) pSNL did not change the number of *Calca* positive neurons in the rostral or middle PBN at 30 dps. (I) pSNL decreased the number of *Cck* positive neurons in the rostral, but not the middle, PBN at 30 dps. (F-I) Rostral sham n = 5, middle sham n = 5, rostral pSNL n = 5, and middle pSNL n = 7. (J) Bilateral injections of AAV1-hSYN-DIO-YFP or AAVDJ-Ef1a-DIO-GFP:TeTx into the PBN of *Calca^Cre/+^* mice. Representative images show expression of YFP and TeTx. Scale bar = 100 µm, dotted line marks the SCP, anterior-posterior Bregma level = -5.1. (K) TeTx expression in PBN *Calca* neurons prevents the development of pSNL driven allodynia measured by von Frey assay. *Calca^Cre/+^*:YFP n = 5 and *Calca^Cre/+^*:TeTx n = 5. (L) Bilateral injections of AAV1-SYN-DIO-mCherry or AAV1-CBA-DIO-hM4Di:mCherry into the PBN of *Calca^Cre/+^* mice. Representative images show expression of mCherry and hM4Di:mCherry. Scale bar = 100 µm, dotted line marks the SCP, anterior-posterior Bregma level = -5.1. (M) hM4Di/CNO inhibition of PBN *Calca* neurons ameliorates pSNL driven allodynia at the 3 dps, measured by von Frey assay. *Calca^Cre/+^*:mCherry n = 5 and *Calca^Cre/+^*:hM4Di n = 5. (N) hM4Di/CNO inhibition of PBN *Calca* neurons ameliorates pSNL driven allodynia at the 30 dps, measured by von Frey assay. *Calca^Cre/+^*:mCherry n = 5 and *Calca^Cre/+^*:hM4Di n = 5.

Inhibition of PBN neurons is known to prevent the manifestation of neuropathic injury-driven allodynia (Zhou et al., 2023; Raver et al., 2020; Sun et al., 2020); however, this phenomenon has never been associated with molecularly defined neurons within the PBN. The PBN *Calca* neurons were an attractive candidate because they are activated by pSNL and many other aversive stimuli (Kang et al., 2022; Campos et al., 2018; Palmiter, 2018). To assess the necessity of *Calca* neurons in the manifestation pSNL-driven chronic pain, we injected an AAV expressing Cre-dependent tetanus toxin light chain (AAV-DIO-GFP:TeTx) bilaterally into the PBN of *Calca^Cre/+^* mice (Fig. 1J) before performing pSNL or sham surgeries. TeTx degrades synaptobrevin, part of the cellular machinery responsible for vesicle release, thus preventing signaling to post-synaptic cells (Kim et al., 2009; Schiavo et al., 2000). Prior to pSNL, a baseline sensitivity to the von Frey (mechanical allodynia) assay was obtained. TeTx expression did not affect baseline paw-withdrawal threshold relative to control animals injected with an AAV expressing Cre-dependent YFP; however, after pSNL, animals treated with TeTx did not develop tactile allodynia (Fig. 1K). Control animals developed allodynia as expected (Fig. 1K). These data suggest that *Calca* neurons are necessary for the manifestation of allodynia in this neuropathic-pain model.

We then asked whether there was a critical window following neuropathic injury during which the activity of *Calca* neurons is necessary for the manifestation of allodynia. To test this possibility, we bilaterally expressed Cre-dependent hM4Di (AAV-DIO-hM4Di:YFP), an inhibitory designer receptor activated by clozapine-N-oxide (CNO) ligand, in the PBN of *Calca^Cre/+^* mice (Fig. 1L). Mice were first assessed at baseline using the von Frey assay; then all animals received unilateral pSNL. The same assay was repeated at 3 dps first following saline injection (0.9%, i.p.), then following CNO injection (5 mg/kg, i.p.). This sequence was performed again at 30 dps. Three days following pSNL, all the mice had developed significant tactile allodynia, CNO-mediated inhibition of *Calca* neurons treatment returned paw withdrawal thresholds to baseline levels (Fig. 1M). At 30 dps, the allodynia was still apparent, but readministering CNO had no effect on pSNL-driven allodynia (Fig. 1N). These data support the hypothesis that PBN *Calca* neurons are necessary for the induction, but not the maintenance, of chronic pain.

### Activation of parabrachial *Calca* neurons is sufficient to drive nociplasticitylII

Chronic stimulation of all excitatory neurons in the PBN can produce a persistent state of allodynia in the absence of tissue injury (Sun et al., 2020), a nociplastic effect. About 85% of PBN neurons are glutamatergic (Pauli et al., 2023; Karthik et al., 2022), including molecularly defined neurons that mediate different and sometimes opposing behavioral effects (Arthurs et al., 2023; Pauli et al., 2023; Karthik et al., 2022; Bowen et al., 2020; Chiang et al., 2020; Chen et al., 2018; Carter et al., 2013). We repeated the experiment of Sun et al. (2020) by bilaterally injecting AAV carrying a Cre-dependent excitatory receptor activated by CNO, (AAV-DIO-hM3Dq:mCherry) into the PBN of *Slc17a6^Cre/+^* mice (*Slc17a6* encodes Vglut2, the vesicular glutamate transporter 2). Daily treatment with CNO (1 mg/kg, i.p., 7 days) produced allodynia that developed following the first CNO injection and lasted many days after the last CNO injection, a sign of nociplasticiy (Fig. S2A). We replicated the results of (Sun et al., 2020), who measured mechanical sensitivity 23 h after each CNO injection; however, when von Frey sensitivity was measured 2 h after each CNO injection it produced a remarkable analgesic effect, that dissipated by 23 h revealing allodynia (Fig. S2A, B). The hM3Dq/CNO-driven analgesia in *Slc17a6^Cre/+^* mice is similar in both scale and transience to the effect of stimulating PBN *Oprm1* neurons, a subset of the greater *Slc17a6* population (Arthurs et al., 2023; Pauli et al., 2023), using the same strategy (Fig. S2C). Consistent with previous studies (Huo et al., 2023), these data show that the PBN *Oprm1* subpopulation can mediate the analgesic effect of stimulating all PBN *Slc17a6* neurons. These findings demonstrate the functional heterogeneity of excitatory neurons within the PBN and are consistent with the molecular heterogeneity of this brain region (Pauli et al., 2023; Karthik et al., 2022).

Given that PBN *Calca* neurons are necessary for the manifestation of neuropathic injury-driven allodynia, we hypothesized that activation of PBN *Calca* neurons might be sufficient to produce nociplasticity. To test this idea, we injected an AAV carrying Cre-dependent hM3Dq:mCherry (or just mCherry as control) bilaterally into the PBN of *Calca^Cre/+^* mice (Fig. 2A). After several weeks for viral expression, 3 consecutive days of CNO delivery (1 mg/kg, i.p.) resulted in significant tactile allodynia 2 h and 23 h post injection that persisted for 10 days following the last CNO injection (Fig. 2B, Fig. S2D). We noted a sexually dimorphic effect in the dissipation, but not the development, of allodynia (Fig. S2E). Additionally, unilateral injection of Cre-dependent hM3Dq:mCherry and subsequent treatment with CNO produced bilateral allodynia (Fig. S2F, G). To determine whether this persistent allodynic effect was in fact nociplasticity, not simply a learned association between the von Frey chamber and *Calca-* neuron-driven aversive sensation, we performed a set of pain assays before and after, but not during, 5 days of CNO administration (Fig. 2C). Two days after the last CNO injection, we observed a significant decrease in von Frey paw-withdrawal threshold, Hargreaves paw-withdrawal latency, and the number of nocifensive behaviors performed on a hot plate (Fig. 2 D-F). The persistent allodynia and hyperalgesia observed in these assays supports our conclusion that chronic activity in the PBN *Calca* population is sufficient to drive nociplasticity.

**Figure 2.**
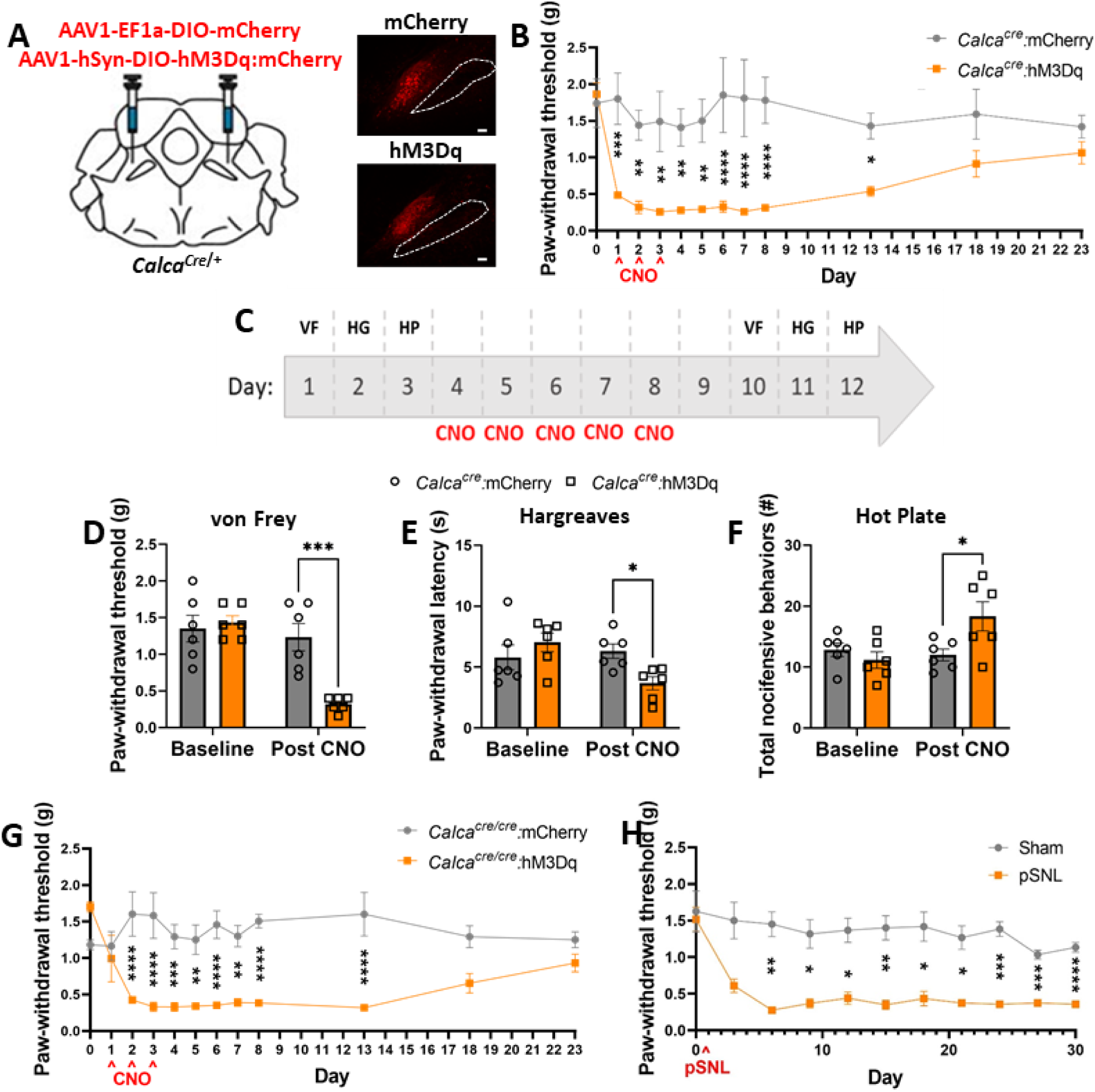
Activation of parabrachial *Calca* neurons is sufficient to drive nociplasticity. (A) Bilateral injections of AAV1-Ef1a-DIO-mCherry or AAV1-hSyn-DIO-hM3Dq:mCherry into the PBN of *Calca^cre/+^* mice. Representative images show expression of mCherry and hM3Dq. Scale bar = 100 µm, dotted line marks the SCP, anterior-posterior Bregma level = -5.1. (B) 3 days of CNO injection (1 mg/kg, i.p.) resulted persistent allodynia measured by von Frey assay. *Calca^cre/+^*:mCherry n = 5 and *Calca^cre/+^*:hM3Dq n = 7. (C) Behavior timeline for von Frey (VF), Hargreave’s (HG), and hot plate (HP) assays before and after 5 consecutive days of CNO injection. *Calca^Cre/+^*:mCherry n = 6 and *Calca^cre/+^*:hM3Dq n = 6. (D) 5 days of CNO injection decreased paw-withdrawal threshold, measured by von Frey assay, which persists after the last CNO injection. (E) 5 days of CNO injection decreased in paw-withdrawal latency, measured by Hargreave’s assay, which persists after the last CNO injection. (F) 5 days of CNO injection increases nocifensive behaviors, measured by hot plate assay, which persists after the last CNO injection. (G) 3 days of CNO injection (1 mg/kg, i.p.) resulted in persistent allodynia measured by von Frey assay. *Calca^Cre/Cre^*:mCherry n = 6 and *Calca^Cre/Cre^*:hM3Dq n = 6. (H) pSNL produced allodynia in *Calca*-null mice measured by von Frey assay.

After observing that activation of PBN *Calca* neurons is both necessary for the manifestation of chronic pain and sufficient to drive nociplasticity, we explored whether expression of the *Calca* gene is a necessary part of these phenomena. For these experiments, we used homozygous *Calca^Cre/Cre^* mice because homozygosity precludes expression of calcitonin gene-related protein, CGRP (Allen et al., 2023; Chen et al., 2018). We performed 3 consecutive days of hM3Dq-mediated stimulation of PBN *Calca^Cre/Cre^* mice, which produced allodynia that lasted 10 days after CNO cessation (Fig. 2G), like the effect observed using heterozygous *Calca^Cre/+^* mice (Fig. 2B). Likewise, pSNL-induced nerve injury in *Calca^Cre/Cre^* mice still produced persistent allodynia (Fig. 2H). In line with previous findings demonstrating that the *Calca* gene is dispensable for acute pain sensation (Zajdel et al., 2021), these data indicate that, while activation of PBN *Calca* neurons is sufficient to drive nociplasticity, the *Calca* gene product, CGRP, does not play a role in this assay. However, it is worth noting that CGRP is important in arthritis-, formalin-, and bladder-pain models (Allen et al., 2023; Shinohara et al., 2017; Han et al., 2015).

### Nociplastic effect scales with the duration of *Calca* neuron activationlII

Seven days of stimulating *Slc17a6* neurons in the PBN produces persistent (>30 days) nociplasticity (Sun et al., 2020 and Fig. S2A) and three days of stimulating *Calca* neurons in the PBN produces a nociplastic effect that lasted 10 days (Fig. 2B). These findings suggest that nociplasticity scales with the duration of *Calca* neuron stimulation. To further examine this phenomenon, we stimulated PBN *Calca* neurons expressing hM3Dq just once and assessed tactile allodynia until paw withdrawal thresholds returned to baseline. One day of CNO-driven stimulation of *Calca* neurons produced 4 days of allodynia (Fig. 3A, B).

**Figure 3.**
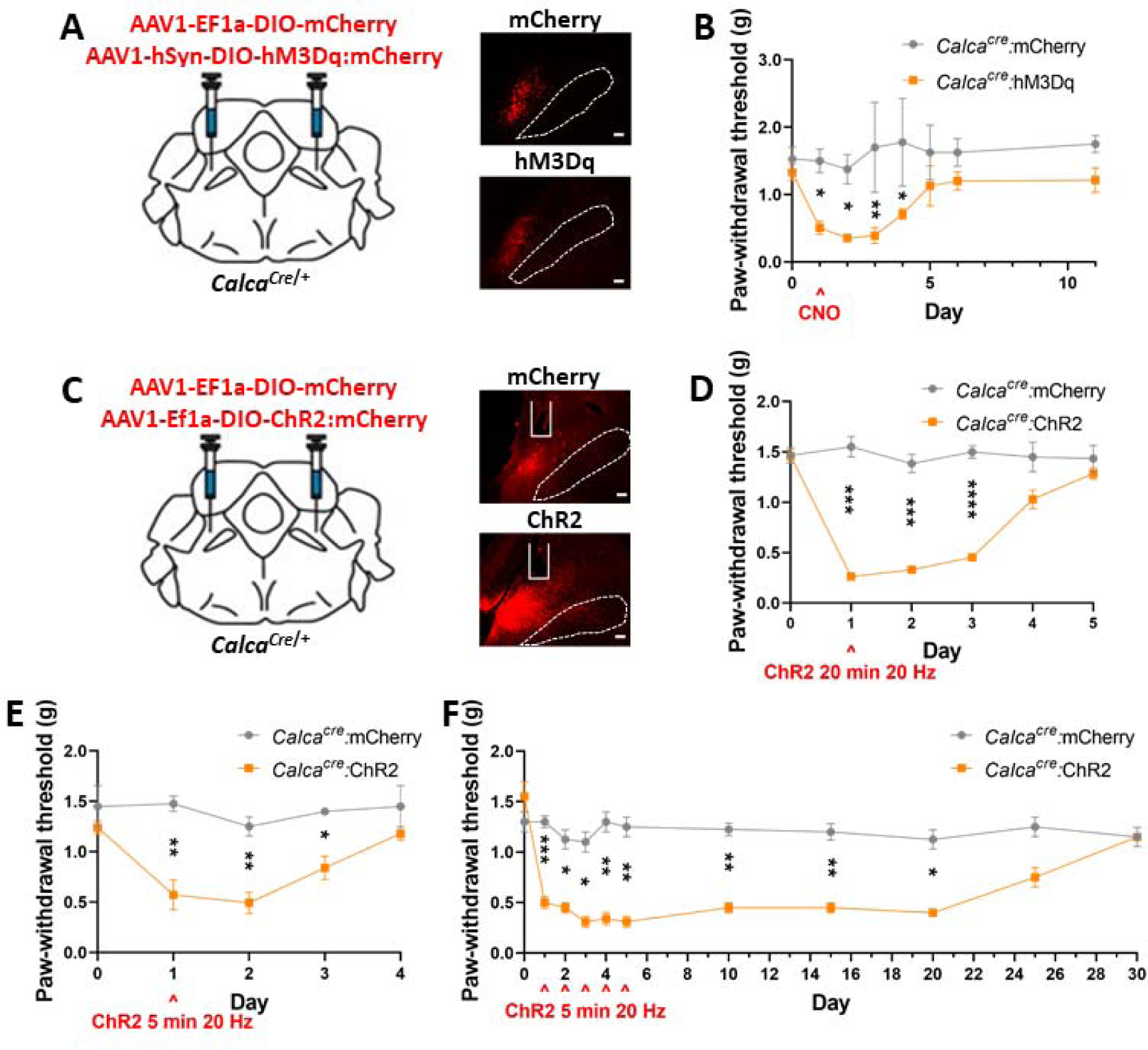
Nociplastic effect scales with the duration of *Calca* neuron activation. (A) Bilateral injections of AAV1-Ef1a-DIO-mCherry or AAV1-hSyn-DIO-hM3Dq:mCherry into the PBN of *Calca^cre/+^* mice. Representative images show expression of mCherry and hM3Dq. Scale bar = 100 µm, dotted line marks the SCP, anterior-posterior Bregma level = -5.1. (B) 1 day of CNO injection (1 mg/kg, i.p.) produced allodynia measured by von Frey assay. *Calca^cre/+^*:mCherry n = 4 and *Calca^cre/+^*:hM3Dq n = 7. (C) Bilateral injections of AAV1-Ef1a-DIO-mCherry or AAV1-Ef1a-DIO-ChR2:mCherry into the PBN of *Calca^cre/+^* mice. Representative images show expression of mCherry and ChR2. Scale bar = 100 µm, dotted line marks the SCP, anterior-posterior Bregma level = -5.1.. (D) 20 min of 473-nm photostimulation (20 Hz, 2 s on 2 s off) resulted in allodynia measured by von Frey assay. *Calca^cre/+^*:mCherry n = 6 and *Calca^cre/+^*:ChR2 n = 7. (E) 5 min of 473-nm photostimulation (20 Hz, 2 s on 2 s off) resulted in allodynia measured by von Frey assay. *Calca^cre/+^*:mCherry n = 4 and *Calca^cre/+^*:ChR2 n = 5. (F) 5 days of 5-min, 473-nm photostimulation (20 Hz, 2 s on 2 s off) resulted in allodynia measured by von Frey assay. *Calca^cre/+^*:mCherry n = 4 and *Calca^cre/+^*:ChR2 n = 4.

Even more acute stimulation was achieved by bilateral stimulation of channelrhodopsin (ChR2) that was targeted to the PBN *Calca* neurons. Twenty min of bilateral 473-nm (20 Hz, 10 mW, 2 s “on”, 2 s “off”) light application over the *Calca* cell bodies produced allodynia within an hour and persisted 2 days after stimulus cessation (Fig. 3C, D). Even 5 min of bilateral stimulation produced allodynia that was weaker, but persisted for 2 days (Fig. 3E). Additionally, 5 min of stimulation performed daily for 5 days produced a nociplastic effect that lasted 15 days after the last stimulation (Fig. 3F). A single 20-min unilateral stimulation using the same parameters also produced bilateral nociplasticity (Fig. S2H). Together these data suggest that the duration of PBN *Calca* neuron stimulation dictates the duration of the resultant nociplasticity.

### Chronic exposure to aversive stimuli drives nociplasticity across a range of of sensory modalitieslII

PBN *Calca* neurons respond to aversive stimuli spanning a range of sensory modalities (Kang et al., 2022; Campos et al., 2018; Chen et al., 2018; Campos et al., 2016; Han et al., 2015). Given that stimulation of *Calca* neurons is sufficient to drive nociplasticity, we explored whether the induction of nociplasticity was agnostic to stimulus modality. Cisplatin chemotherapy (Alhadeff et al., 2017; Park et al., 2013; Ta et al., 2009), lithium chloride (LiCl)-induced visceral malaise (Chen et al., 2018; Carter et al., 2015), foot shock (Campos et al., 2018), and the threat of predation (Kang et al., 2022) have all be demonstrated to activate PBN *Calca* neurons. Chronic exposure to cisplatin (Park et al., 2013), nitroglycerin (NTG; a model for migraine pain) (Pradhan et al., 2014), and foot shock (Wu et al., 2020) are known to produce persistent allodynia. Consistent with previous studies (Pradhan et al., 2014), we found that 5 days of NTG exposure (10 mg/kg, i.p.) produced persistent allodynia (Fig. 4A). We also found that 3 days of injection with LiCl (0.2 M at 15 mL/kg, i.p.) produced persistent allodynia (Fig. 4B). Even 3 days of exposure to a predatory threat (5-min chase daily for 3 days with a toy robotic bug) could promote mild nociplasticity, measured by von Frey assay (Fig. 4C).

**Figure 4.**
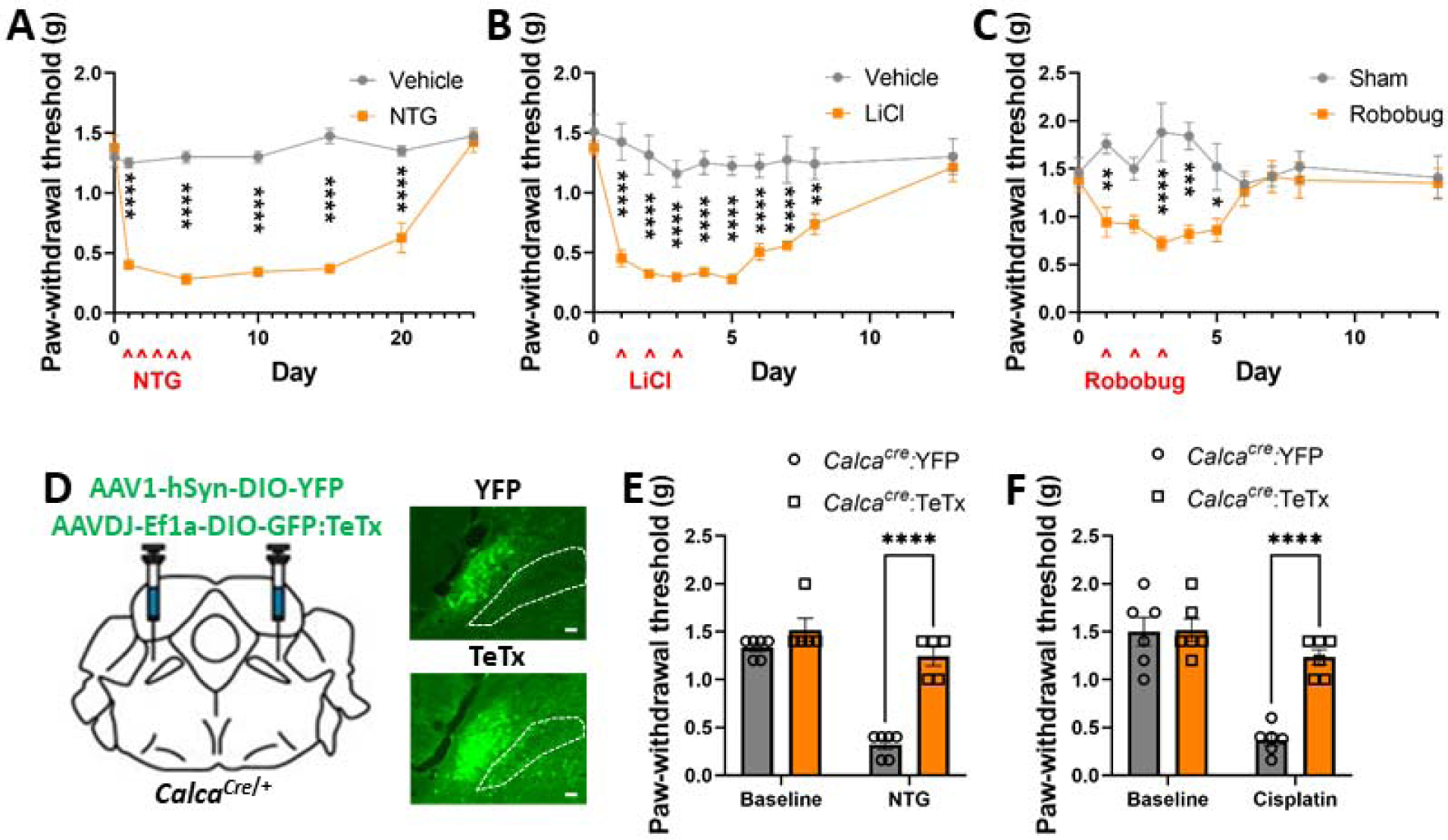
Chronic exposure to all aversive stimuli tested drives nociplasticity regardless of sensory modality. (A) 5 days of NTG exposure (10 mg/kg, i.p.) produced allodynia measured by von Frey assay. Vehicle n = 4 and NTG n = 4. (B) 3 days of LiCl exposure (0.2 M, 15 mL/kg, i.p.) produced allodynia measured by von Frey assay. Vehicle n = 6 and LiCl n = 8. (C) 3 days of robobug chase (10 min) produced allodynia measured by von Frey assay. Vehicle n = 5 and robobug n = 5. (D) Bilateral injections of AAV1-hSYN-DIO-YFP or AAVDJ-Ef1a-DIO-GFP:TeTx into the PBN of *Calca^Cre/+^* mice. Representative images show expression of YFP and TeTx. Scale bar = 100 µm, dotted line marks the SCP, anterior-posterior Bregma level = -5.1. (E) TeTx expression in PBN *Calca* neurons prevented the development of NTG-driven allodynia measured by von Frey assay. *Calca^Cre/+^*:YFP n = 6 and *Calca^Cre/+^*:TeTx n = 5. (F) TeTx expression in PBN *Calca* neurons prevented the development of cisplatin-driven allodynia measured by von Frey assay. *Calca^Cre/+^*:YFP n = 6 and *Calca^Cre/+^*:TeTx n = 6.

NTG- and cisplatin-produced nociplasticity were prevented by prior expression of TeTx in *Calca* neurons (Fig. 4D, E, F). Interestingly, the photophobia that developed following NTG injection did not persist as long as the allodynia (Fig. S2I). Additionally, despite pretreatment of the PBN *Calca* neurons with TeTx, photophobia still occurred following NTG injection (Fig. S2J). These data suggest that chronic exposure to these aversive stimuli, spanning a range of sensory modalities, is capable of inducing nociplasticity via activation of *Calca* neurons.

### *Calca* neurons exhibit plasticity following activation

We have established that *Calca* neuron activity is sufficient to drive nociplasticity; however, the neuronal populations exhibiting neuroplastic changes are not yet established. We performed patch-clamp electrophysiology to assess the excitability of *Calca* neurons expressing hM3Dq:mCherry following 3 days of either CNO or saline injections (Fig. 5A, B, C). Forty-eight hours following the final injection, at which point all injected CNO should be metabolized (Raper et al., 2017), *Calca* neurons in slices were injected with 800-ms pulses of current ranging from 0 to 240 pA (Fig. 5C, D). Quantification of the elicited spikes revealed two distinct response patterns, regular and late firing (Fig. 5D, F). Both the regular- and late-firing populations were significantly more responsive to current injection when the mice were pretreated with CNO rather than saline (Fig. 5E, G). These data demonstrate that chronic activation of *Calca* neurons produces a persistent increase in their intrinsic excitability.

**Figure 5.**
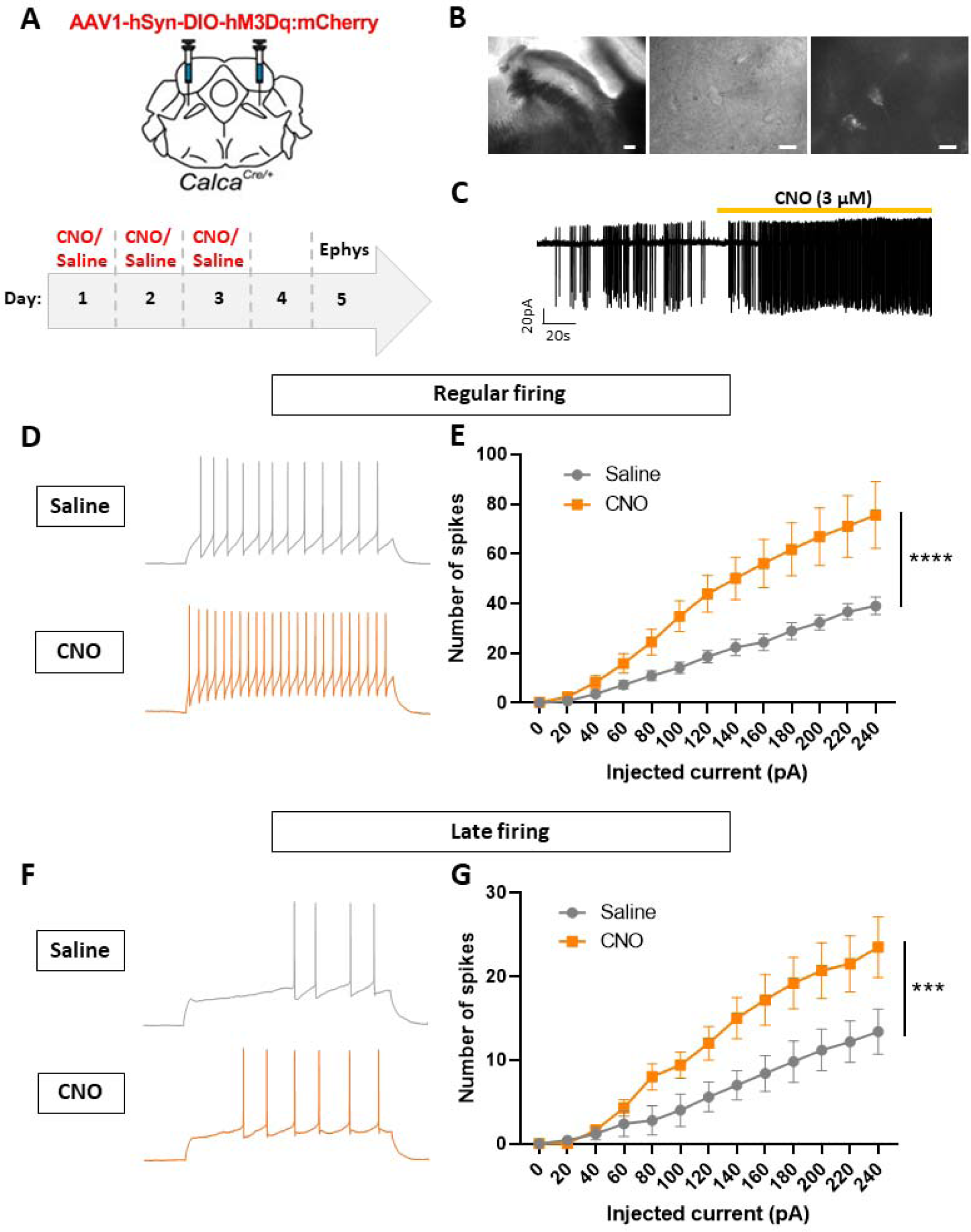
Chronic stimulation of PBN *Calca* neurons alters their intrinsic excitability. (A) Bilateral injections of AAV1-Ef1a-DIO-mCherry or AAV1-hSyn-DIO-hM3Dq:mCherry into the PBN of *Calca^cre/+^* mice followed by 3 days of CNO or saline injection and 48 h of no stimulation prior to electrophysiology. (B) Representative images of PBN (10x; scale bar, 100 µm), patched cell (40x; scale bar, 10 µm), and hM3Dq expression (40x; scale bar, 10 µm) under patch microscope. (C) Representative trace of spikes elicited by CNO. (D) Representative traces showing regular firing *Calca* neurons. (E) 3 days of CNO injection (i.p.) prior to electrophysiology resulted in an increase in the number of spikes elicited by current injection in the regular-firing population. Saline-treated n = 3 animals, 14 neurons; CNO treated n = 3 animals, 11 neurons. (F) Representative traces showing late firing *Calca* neurons. (G) 3 days of CNO injection (i.p.) prior to electrophysiology resulted in an increase in the number of spikes elicited by current injection in the late-firing population. Saline-treated n = 3 animals, 5 neurons; CNO treated n = 3 animals, 11 neurons.

Because NTG administration induces *Calca*-neuron-dependent persistent allodynia, like the effect of hM3Dq/CNO mediated stimulation, we hypothesized it does so by increasing the activity of *Calca* neurons. To investigate this hypothesis, we conducted Ca^2+^ imaging experiments in which we tracked the activity of individual *Calca* neurons across multiple days. To monitor the activity of *Calca* neurons in behaving animals AAV-DIO-GCaMP6m was expressed, and Gradient Refractive Index (GRIN) lens was implanted over the PBN of *Calca^Cre/+^* mice. After 4 weeks of recovery, a microendoscope was attached to each mouse and left in place for 4 consecutive days. This strategy ensured a stable field of view for tracking individual neurons each day. On day 1, mice received an intraperitoneal injection of vehicle and on day 2 they received NTG (10 mg/kg, i.p.). After each injection, mice were placed in the von Frey-testing apparatus for 35 min of acclimation, followed by 5 min of baseline neuronal activity measurement, and then 10 min of von Frey testing (8 filament applications, with a 1-min intertrial interval). On days 3 and 4, *Calca* neurons were imaged again without further injections to determine if NTG treatment had residual effects that could drive persistent allodynia (Fig. 6A).

**Figure 6.**
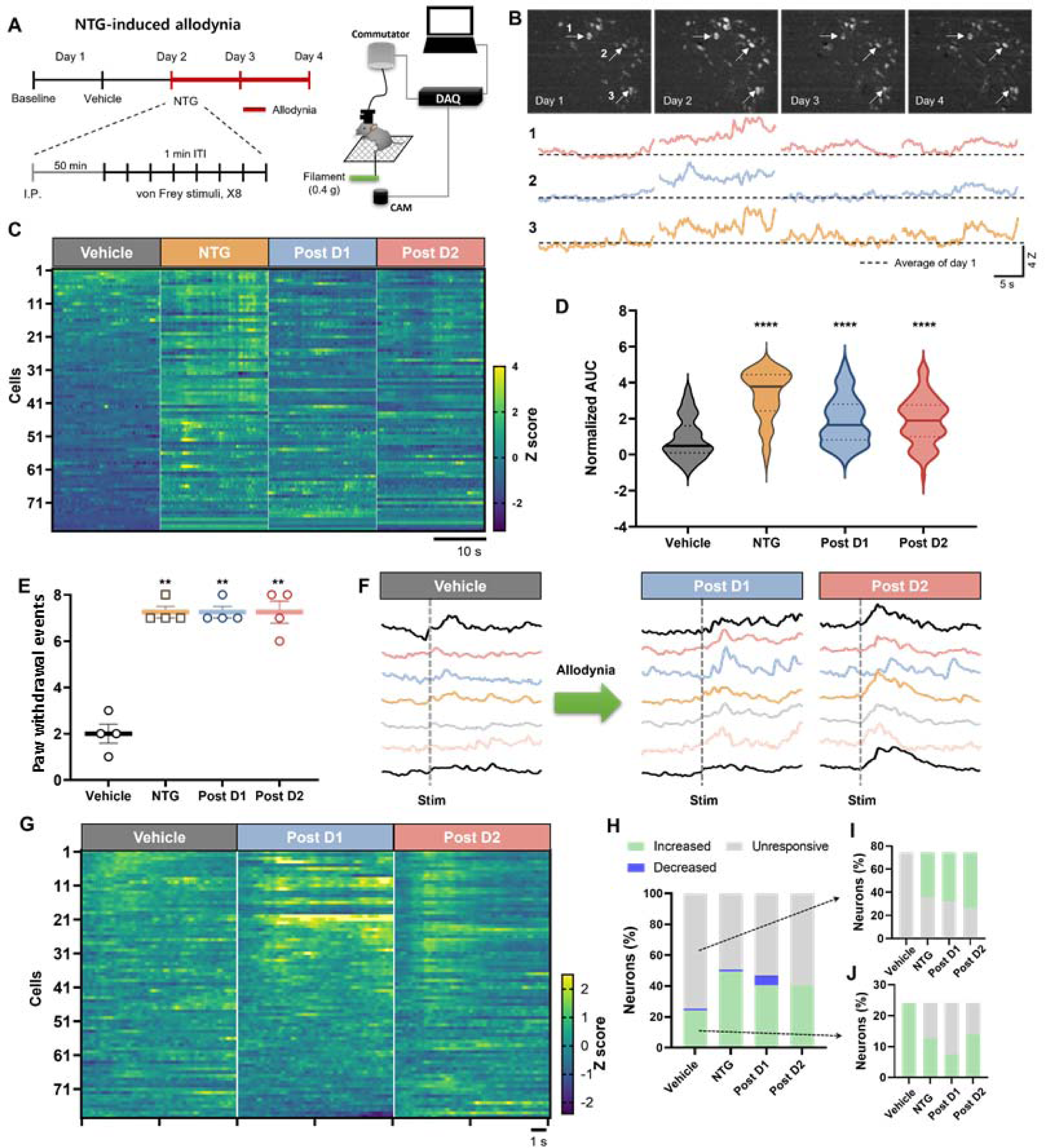
*Calca* neuron activity during the development of mechanical allodynia. (A) Schematic depiction of Ca2+ imaging during NTG induced mechanical allodynia. (B) Multi-day tracking of *Calca* neuron fluorescent activity. Top; Field of view (FOV) of representative animal for 4 days. Bottom; representative neural activities throughout 4 days of 3 neurons marked by arrows in FOVs. (C) NTG injection increased unstimulated neuronal activity. Elevated neuronal activity persisted 24 and 48 h post injection. Individual neurons are aligned across days in heatmap. (D) Average calcium transient area under the curve increased following NTG injection. This increase in fluorescent activity remained elevated 24 and 48 h post injection. (E) NTG injection increased the number of times out of 8 applications that mice responded to a 0.4 g von Frey filament. Bar indicates mean ± S.E.M. (F) Representative traces of filament-evoked neural activities. (G) The number of *Calca* neurons responsive to application of a 0.4 g von Frey filament increased after NTG injection. The increase in responsive neurons persisted 24 and 48 h after NTG injection. (H) The percentage of *Calca* neurons responsive to application of a 0.4 g von Frey filament increased from 24% after vehicle injection to 49.5% after NTG injection. The percent of 0.4 g von Frey filament-responsive neurons remained elevated, at 40.5%, 24 and 48 h after NTG injection. (I) The majority of neurons unresponsive to 0.4 g von Frey filament application following vehicle injection (i.p.) became responsive to the 0.4 g filament following NTG injection (i.p.). (J) About half of neurons responsive to 0.4 g von Frey filament application following vehicle injection (i.p.) became unresponsive to the 0.4 g filament following NTG injection (i.p.).

We tracked the same 79 neurons throughout the experiment (Fig. 6B). NTG administration increased the basal fluorescence level of individual neurons dramatically compared to vehicle administration (Fig. 6B, C). Notably, these elevated fluorescent activities were sustained on days 3 and 4 following NTG injection (Fig. 6B, C); 60% of neurons continued to exhibit elevated basal fluorescence on days 3 and 4 (Fig. 6C, Fig. S3A). The number and amplitude of Ca^2+^ transients did not change across days (Fig. S3B, C). The averaged basal fluorescence level of all 79 neurons (4 mice) increased after NTG administration and remained elevated for two days (Fig. 6D).

We also examined whether NTG administration was accompanied by an increase in the von Frey-elicited responses of individual neurons. NTG injection increased paw-withdrawal responses of all the mice to a low threshold (0.4 g) von Frey filament (Fig. 6E). Some neurons that exhibited increased activity following von Frey stimuli to vehicle injection showed similar activity after NTG injection (Fig. 6F), while most *Calca* neurons displayed increased fluorescence only after NTG injection (Fig. 6F, G). On the vehicle-injection day, 24% of neurons were activated by the 0.4-g von Frey filament, whereas 49.4% were activated after NTG treatment and 40.5 % responded on days 3 and 4. (Fig. 6H). Analysis of individual *Calca* neurons across experimental days revealed that 56.7% of the initially unresponsive neurons became responsive after NTG, with similar values on days 3 and 4 (Fig. 6I). Of the von Frey-responsive neurons on the vehicle-injection day, about half lost their responsiveness following NTG injection and across the subsequent imaging sessions (Fig. 6J). There were no persistent changes in the area under the curve of calcium transients, the number of calcium transients, or the amplitude of calcium transients following 0.4-g von Frey stimulation (Fig. S3D, E, F). These data suggest that the primary effect of NTG treatment is the recruitment of more *Calca*-responsive neurons rather than an increase in the frequency or magnitude of responses.

### Intrathecal injection of neuropeptide Y does not reverse hM3Dq/CNO-driven allodynia

How does manipulation of neurons in the brain result in tactile allodynia allowing gentle touch to appear painful? The allodynia that develops after sciatic nerve injury has been shown to involve plasticity in spinal dorsal horn inhibitory neurons that normally prevent (gate) low-threshold primary afferent (Aß) activity from reaching the nociceptive spinoparabrachial projection neurons, thereby allowing gentle touch to the hind paw to promote paw withdrawal as if it was painful (Cao et al., 2022; Nelson et al., 2022; Peirs et al., 2021; Boyle et al., 2019; Petitjean et al., 2015; Lu et al., 2013; Todd et al., 2010). We hypothesized that chemogenetic activation of *Calca* neurons may activate descending circuits to the spinal cord resulting in plasticity that resembles that induced by nerve injury. Pharmacological or genetic manipulations of several different populations of excitatory neurons in the spinal cord have been shown to reverse peripheral nerve injury-induced allodynia (Cao et al., 2022; Nelson et al., 2022; Peirs et al., 2021). For example, intrathecal injection of the NPY Y1 receptor 1 agonist NPY^Leu,Pro^ has been shown to transiently reduce allodynia after sciatic nerve injury (Nelson et al., 2022, Nelson et al., 2021; Malet et al., 2017). Thus, we tested our hypothesis by intrathecal injection of NPY^Leu,Pro^ into mice with pSNL (as positive control) or mice that had developed tactile allodynia after 3 days of *Calca* neuron activation (Fig. 7A). NPY^Leu,Pro^ injections ameliorated the allodynia produced by pSNL as expected but had no effect on the allodynia that developed after *Calca* neuron activation (Fig. 7B), indicating that the nociplastic allodynia that develops after *Calca* neuron activation does not rely on the same mechanisms as nociplasticity driven by pSNL.

**Figure 7.**
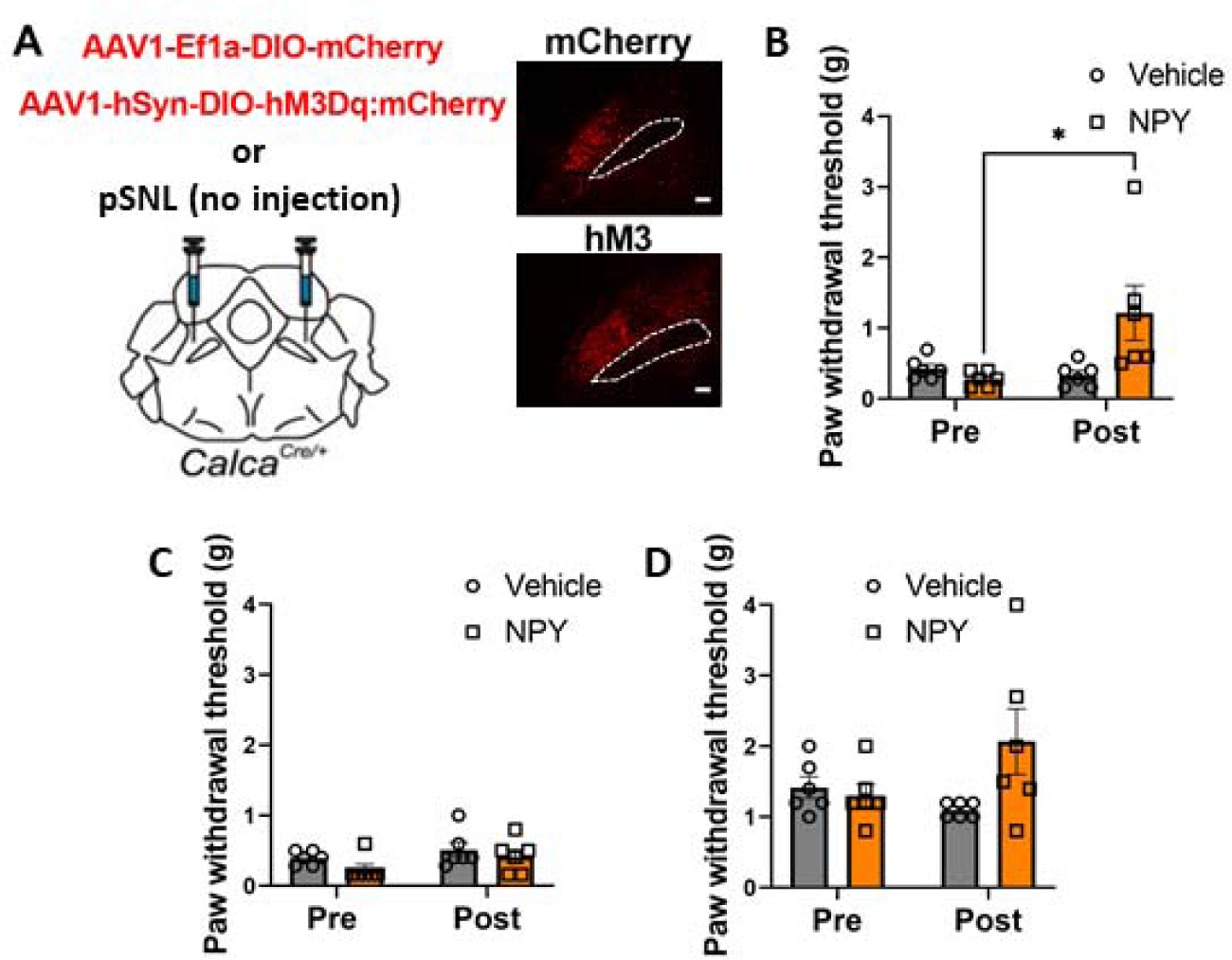
Intrathecal NPY does not reverse hM3Dq/CNO-driven allodynia. (A – J) n = 4 animals, 79 neurons. (A) pSNL or bilateral injections of AAV1-Ef1a-DIO-mCherry or AAV1-hSyn-DIO-hM3Dq:mCherry into the PBN of *Calca^cre/+^* mice. Representative images show expression of mCherry and hM3Dq. Scale bar = 100 µm, dotted line marks the SCP, anterior-posterior Bregma level = -5.1. (B) Intrathecal (i.t.) injection of NPY^Leu,Pro^ into pSNL animals reversed pSNL driven allodynia. (C) I.t. injection of NPY^Leu,Pro^ into Calca^Cre^:hM3Dq animals treated with CNO for 3 days did not reverse allodynia. (C) I.t. injection of NPY^Leu,Pro^ into control Calca^Cre^:mCherry animals treated with CNO for 3 days did not affect the paw withdrawal threshold.

## DISCUSSION

We demonstrate that *Calca-*neuron activity is necessary for the induction of pSNL-induced allodynia, and that daily chemogenetic or optogenetic stimulation of *Calca* neurons is sufficient to induce allodynia and hyperalgesia that persists for many days beyond the stimulation period. These chronic pain symptoms are the result of a generalized nociplasticity and not a learned association between the assays and the aversion generated by *Calca* neuron activity. Although *Calca* neuron activity is both necessary and sufficient to generate chronic pain, *Calca* gene products (i.e., CGRP) themselves did not play a significant role in this effect, which is unexpected because CGRP signaling has been shown to be important in other pain models (Allen et al., 2023; Shinohara et al., 2017; Han et al., 2015).

Sun et al., (2020) reported that 7 days of chemogenetic activation of all glutamatergic neurons in the PBN could produce allodynia that lasted for many days after the final CNO injection. We obtained similar results when assaying von Frey sensitivity 23 h after each CNO injection, when CNO would have been cleared from the circulation. Remarkably, when we assayed allodynia 2 h after each CNO injection, when hM3Dq would still be activated, there was a dramatic analgesic effect. A similar effect was achieved with hM3Dq-mediated stimulation of *Oprm1* neurons. In contrast, chemogenetic activation *Calca* neurons, which represent ∼15% of the glutamatergic neurons (Pauli et al., 2022), drives allodynia at both 2 and 23 h after CNO. This result suggests that the glutamatergic population includes neurons that promote allodynia (*Calca* neurons) and those that promote analgesia (*Oprm1* neurons). Both genes are expressed in several molecularly defined clusters in the PBN (Pauli et al., 2022). The activation of *Oprm1* neurons is aversive (Liu et al., 2022), so it is likely the analgesia we observed was “stress-induced” (Butler and Finn, 2009).

*Calca-*neuron-mediated nociplasticity scales with stimulus duration. Multiple rounds of chemogenetic stimulation produced allodynia that persisted longer than a single stimulation. Likewise, a brief optogenetic stimulation produced allodynia that was less persistent than after multiple stimulations. The stimulus scaling of nociplasticity that we observed is consistent with the conclusion that *Calca* neuron activity drives nociplasticty.

Several studies, ours included, have demonstrated that unilateral neuropathic injury can produce bilateral pain (Sugimoto et al., 2021; Abraham et al., 2020; Raver et al., 2020; Arguis et al., 2008; Koltzenberg et al., 1999). This phenomenon indicates that persistent pain has the capacity to induce generalized nociplasticity. Because artificial activation of *Calca* neurons is sufficient to produce generalized pain and that a wide variety of aversive situations activate *Calca* neurons, some of which are not generally considered to be painful, we predicted that many aversive experiences, especially if repeated, would have the capacity to drive nociplasticity. Chronic exposure to NTG (a model of migraine) was shown to produce persistent allodynia (Pradhan et al., 2014), an effect that we duplicated and went on to show was dependent on activation of *Calca* neurons. Similarly, cisplatin-induced chemotherapy can produce allodynia (Park et al, 2013), activate *Calca* neurons (Ta et al., 2009; Park et al., 2013; Alhadeff et al., 2017) and produce persistent allodynia that we show depends on *Calca*-neuron activation in the PBN. We also demonstrate that prolonged visceral malaise (nausea) induced by treatment with LiCl, as well as predatory simulation are sufficient to induce allodynia that may persist for many days. These findings point to a mechanism by which diverse and polymodal aversive experiences can produce nociplastic pain phenotypes via the activation of PBN *Calca* neurons. An implication of this result is that nociplastic pain may be produced by adverse life events that are distinct from somatic nerve injury. This phenomenon has been observed epidemiologically. Individuals with adverse childhood experiences are significantly more likely to develop chronic pain (Groenewald et al., 2020; Dokyoung et al., 2019; Sherman et al., 2015). These findings may also explain the high degree of mixed pain pathophysiology in chronic pain patients (Fitzcharles et al., 2021). In this circumstance, it is possible that persistent nociceptive or neuropathic pain drives nociplastic pain via the chronic activation of *Calca* neurons.

*Calca* neurons exhibit plasticity in the form of increased intrinsic excitability following three days of CNO stimulation of hM3Dq and an increase in the number of neurons responsive to innocuous tactile stimulation for several days after NTG exposure. These effects may initiate a persistent pain state, but our data showing *Calca* neuron activity at 3 days but not 30 days, and hM4Di inhibition data demonstrating reversal of allodynia at 3 days but not 30 days, suggest that *Calca* neurons are no longer active 30 days after pSNL when allodynia persists. We suggest that long-lasting allodynia involves nociplastic changes in the circuitry downstream of *Calca* neurons (Qi et al., 2022; Zhou et al., 2023; Cai et al., 2014; Wilson et al., 2019; Chen et al., 2017) possibly including the spinal microcircuitry. Chronic pain after sciatic nerve injury changes the microcircuitry of the dorsal horn, allowing Aβ fibers to indirectly stimulate spinoparabrachial projection neurons via a neural circuit that is inhibited under non-painful conditions (Cao et al., 2022; Nelson et al., 2022; Peirs et al., 2021; Boyle et al., 2019; Petitjean et al., 2015; Lu et al., 2013; Todd et al., 2010). Intrathecal injection of neuropeptide Y (NPY) can reverse neuropathic pain (Nelson et al., 2022; Nelson et al., 2021), a result that we confirmed in the pSNL model. However, NPY did not ameliorate the tactile allodynia that develops after hM3Dq/CNO activation of *Calca* neurons, suggesting that neuroplasticity that promotes allodynia after pSNL is distinct from that that occurs after *Calca*-mediated allodynia.

In conclusion, our study highlights the critical role of *Calca* neurons in driving nociplasticity and persistent pain states. We have shown that activation of *Calca* neurons induces allodynia and hyperalgesia that persist beyond the stimulation period, implicating a generalized nociplasticity rather than learned associations. Furthermore, our findings suggest that diverse aversive experiences can activate *Calca* neurons and contribute to the development of nociplastic pain, shedding light on the potential mechanisms underlying chronic pain conditions that extend beyond somatic nerve injury. This research underscores the complexity of pain pathophysiology and offers new insights into the interplay between neural circuits and aversive life events in shaping chronic-pain phenotypes.

## STAR*METHODS

Detailed methods are provided in the online version of this paper and include the following:

- KEY RESOURCES TABLE
- RESOURCE AVAILABILITY
  ○ Lead contact
  ○ Materials availability
  ○ Data and code availability
- EXPERIMENTAL MODEL AND STUDY PARTICIPANTS
  ○ Mice
- METHOD DETAILS
  ○ Virus production
  ○ Partial sciatic nerve ligation
  ○ Intrathecal injection of NPY
  ○ Stereotaxic surgery
  ○ Allodynia assays
  ○ Pharmacological injections
  ○ Optogenetic stimulation
  ○ Immunohistochemistry
  ○ RNAscope *in situ* hybridization
  ○ Electrophysiology
  ○ Calcium imaging
- QUANTIFICATION AND STATISTICAL ANALYSIS

## KEY RESOURCES TABLE

| Reagent | Source | Identifier |
| --- | --- | --- |
| **Antibodies and RNAscope probes** |  |  |
| Rabbit-anti-dsRed | Takara | ab 632496 |
| Chicken-anti-GFP | Abcam | ab 13970 |
| Alexa Fluor 594 donkey anti-rabbit | Jackson ImmunoResearch | Cat #715-585-150, RRID: AB_2340854 |
| Alexa Fluor 488 donkey anti-chicken | Jackson ImmunoResearch | Cat #703-545-155, RRID: AB_2340375 |
| RNAscope Probe Mm-*Fos*-C1 | ACD Biotechne | Ref #316921 |
| RNAscope Probe Mm-*Calca*-C2 | ACD Biotechne | Ref #578771-C2 |
| RNAscope Probe Mm-*Cck*-C2 | ACD Biotechne | Ref #402271-C3 |
| **Virus strains** |  |  |
| pAAV1-SYN-DIO-YFP | Karl Diesseroth | Similar to Addgene plasmid #27056 |
| pAAVDJ-Ef1a-DIO-GFP:TeTx | Janelia | N/A |
| pAAV1-Ef1a-DIO-ChR2:mCherry | Karl Diesseroth | Addgene #35508 |
| pAAVDJ-SYN-DIO-hM3Dq:mCherry | Bryan Roth | Addgene #44361 |
| pAAV1-Ef1a-DIO-mCherry | Bryan Roth | Addgene #50462 |
| pAAV1-Ef1a-DIO-GCaMP6m | Larry Zweifel | N/A |
| pAAV1-hSyn-DIO-hM4:mCherry | Bryan Roth | Addgene #44362 |
| **Chemicals** |  |  |
| Clozapine-N-Oxide | Fisher Scientific | Cat #L121, CAS 7447-41-8 |
| Nitroglycerin | American Regent, Inc. | NDC 0517-4810-25 |
| Lithium Chloride | Fisher Scientific | Cat #L121, CAS 7447-41-8 |
| Cisplatin | West-Ward | NDC 0143-9504-01 |
| Paraformaldehyde 32% Aqueous Solution | Electron Microscopy Sciences | Cat #15714-S |
| Phenobarbitol | Akorn | NDC 59399-185-90 |
| Normal donkey serum | Jackson ImmunoResearch | Cat #017-000-121, RRID: AB_2337258 |
| O.C.T. | Thermofisher | Cat #23-730-571 |
| DAPI Fluoromount-G | SouthernBiotechne | Cat #0100-20 |
| [Leu<sup>31</sup>, Pro<sup>34</sup>]-NPY | Tocris | Cat #1176 |
| **Experimental Models** |  |  |
| Mouse: C57BL/6J | Jackson Laboratory | JAX:000664 |
| Mouse: *Calca*<sup>Cre</sup> | Jackson Laboratory | JAX:033168 |
| Mouse: *Slc17a6*<sup>Cre</sup> | Jackson Laboratory | JAX:016963 |
| Mouse: *Oprm1*<sup>Cre</sup> | Jackson Laboratory | JAX:035574 |
| **Software and algorithms** |  |  |
| QuPath | GitHub | https://qupath.github.io/ |
| Prism (v9.5.1) | GraphPad Software | www.graphpad.com |
| Fiji/ImageJ | Fiji/NIH | www.fiji.sc |
| pClamp 11 | Molecular Devices | www.moleculardevices.com |
| Clampfit 11.0.3 | Molecular Devices | www.moleculardevices.com |
| IDAS | Inscopix | www.inscopix.com |
| IDPS | Inscopix | www.inscopix.com |
| **Hardware** |  |  |
| Modular chambers and elevated mesh bottom stand | Bioseb | Ref #BIO-PVF |
| Plantar test for thermal stimulation | Ugo Basil | Product code: 37370 |
| Cold hot plate | Bioseb | Ref #BIO-CHP |

## RESOURCE AVAILABILITY

### Lead contact

Further information and request for resources and reagents should be directed to and will be fulfilled by Richard Palmiter, the lead contact.

### Materials availability

This study did not generate any new reagents or behavioral systems.

### Data and code availability

- Data will be made available upon reasonable request to the corresponding author.
- All original code has been deposited at GitHub and is publicly available.
- Any additional information required to reanalyze the data reported in the manuscript is available from the lead contact upon request.

## EXPERIMENTAL MODEL AND STUDY PARTICIPANT DETAILS

### Mice

All experiments followed protocols approved by the Institutional Animal Care and Use Committee at the University of Washington and were in accordance with the National Institute of Health guidelines for animal research. Experiments on wild-type animals used C57BL/6 mice. Most experiments used heterozygous *Calca^Cre/+^* or homozygous *Calca^Cre/Cre^* mice on a C57BL/6 genetic background were generated and maintained as described (Carter et al., 2013). One set of experiments used *Slc17a6^Cre/+^* or *Oprm1^Cre/+^* mice. Male and female animals were 7 to 9 weeks of age at the onset of all experiments and no more than 18 weeks by their experimental endpoint. Prior to surgical manipulation animals were group housed, had *ad libitum* access to food and water, and were kept on a 12-h, light-dark cycle at 22°C. After surgical manipulation animals were singly housed, all other housing parameters remained consistent. All experiments included 4-10 experimental animals (e.g., transduced with viruses allowing Cre-dependent expression of hM3Dq, hM4Di, ChR2, TeTx) and 4-7 control animals (e.g., transduced with mCherry or YFP) to account for potential variation between experimental sessions. Animals from the same litter were randomly assigned to experimental or control groups. All experiments were performed blind, except the *Oprm1^Cre/+^* experiment. Individual cohorts of mice were used for all experiments except those mentioned below. The same cohort of *Calca^Cre/+^*:TeTx animals was used in the NTG and cisplatin experiments with 1 week of recovery between NTG and cisplatin injections. The same cohort of *Calca^Cre/+^*:ChR2 animals was used for all ChR2 experiments with 1 week of recovery between the 1-day ChR2 stimulation and the 5-day ChR2 stimulation experiment. Viral transduction was assessed at the end of each experiment histologically and only mice with correct expression were included in data analysis.

## METHOD DETAILS

### Virus production

AAV1-Ef1a-DIO-mCherry and AAVDJ-SYN-DIO-hM3Dq:mCherry DNA plasmids were provided by B. Roth (Addgene #50462 and #44361). AAV1-SYN-DIO-YFP and AAV1-Ef1a-DIO-ChR2:mCherry DNA plasmids were provided by K. Diesseroth (Stanford University). AAVDJ-Ef1a-DIO-GFP:TeTx and AAV1-CBA-DIO-hM4Di:YFP plasmids were constructed by R. Palmiter (University of Washington). AAV1-CBA-DIO-GCaMP6m DNA plasmid was provided by L. Zweifel. AAV1 serotype viruses were prepared in-house by transfecting HEK cells with each of these plasmids. Viruses were purified by sucrose and CsCl gradient centrifugation steps and re-suspended in 0.1 M phosphate-buffered saline (PBS) at about 10^13^ viral particles/mL. AAVDJ serotype viruses were prepared by Janelia Viral Tools lab.

### Partial sciatic nerve ligation

Partial sciatic nerve ligation (pSNL) was performed as described (Abaraham et al., 2020). In brief, mice were anesthetized with 2% isoflurane at a flow rate of 1 L/min. Subsequently a 2-cm incision was made over the lateral aspect of the proximal third of the left hind leg. Blunt dissection was used to visualize the sciatic nerve, which was then exteriorized with hooked forceps. For sham surgeries the sciatic nerve was then immediately returned to its position. For nerve ligation surgeries, a 6-0 silk suture was passed through 30-50% of the nerve bundle, before being tightly ligated and crushed. The skin was then closed with 6-0 silk sutures.

### Intrathecal injection of NPY

Intrathecal injection of [Leu^31^, Pro^34^]-NPY (TOCRIS) was performed as described by Nelson et al. (2022). In brief, a 30G needle attached to a 25 µL microsyringe (Hamilton) was inserted between the L5/L6 vertebrae, puncturing the dura mater; 5 µL of vehicle (0.9 % saline) or NPY was injected. Each mouse was injected twice, once with vehicle and once with NPY, with 48 h between injections.

### Stereotaxic surgery

Mice were anesthetized with 2% isoflurane at a flow rate of 1 L/minute. After anesthesia induction mice were placed on a stereotaxic frame (David Kopf Instruments). Stereotaxic coordinates in the anterior posterior plane were normalized using a correction factor (F = (Bregma – Lambda)/4.21). Viral injections were performed bilaterally for all experiments into the PBN (anterior-posterior: -1.3, rostral-caudal: -4.8, medial-lateral: +/-1.3) at a rate of 0.2 µL/min for 2.5 min for a total of 0.5 µL. At the end of each experiment animals were euthanized with phenobarbital and the brain was extracted for histological evaluation of viral placement and fiber optic placement.

For *in vivo* Ca^2+^ imaging experiments, viral injections were in the external lateral part of the parabrachial nucleus (PBel) using the following coordinates relative to bregma at the skull surface: AP -4.90 mm; ML ±1.35 mm; DV +3.40. Viruses were injected unilaterally (randomly assigned) 0.5 μl at 0.1 ml/min. A gradient refractive index (GRIN) lens was positioned above the target area (AP -4.80 mm; ML ±1.7 mm; DV +3.65), and three tungsten wires protruding approximately 0.5 mm beyond the lens surface (A-M systems) were attached to the lens to reduce motion artifacts during imaging. The lens was then lowered at a rate of 0.1 mm/min. To secure the lens to the skull, super glue (C&B Metabond from Parkell) and dental cement were used. After 4 weeks of recovery, mice were tested to ensure field of view (FOV). Animals with a stable FOV were used in the experiments for the subsequent 2-4 weeks.

### Allodynia assays

The von Frey, tactile-sensitivity assay was performed using the ascending application method as described (Abraham et al., 2020). Each animal was placed in a 11.5-cm by 7.5-cm chamber with a wire mesh floor (Bioseb). Animals were acclimated to the chamber for 30 min before the assay began. Filaments were applied to the plantar surface of each hind paw a total of 5 times. Filament application began with a 0.16-g filament and ended after two consecutive filaments elicit a paw withdrawal on 3 or more out of 5 applications. The paw-withdrawal thresholds for each hind paw were measured and averaged because there was no significant difference in the left/right paw-withdrawal thresholds following any of the performed manipulations (pSNL, NTG, LiCl, etc.).

The Hargreaves thermal-sensitivity assay was performed as described (Wilson et al., 2019). Each animal was placed in an 11.5-cm by 7.5-cm chamber with a wire mesh floor. Animals were acclimated to the chamber for 30 min before the assay began. Each animal received infrared thermal stimulation (Ugo Basil, model 37370) a total of 3 times on both hind paw plantar surfaces. Latency to paw withdrawal was averaged across the 3 sessions. The paw-withdrawal latency for each paw was averaged as we did not see a difference in their response latency values.

The hot-plate assay was performed as described (Nelson et al., 2022). Each animal was placed in a 16.5-cm by 16.5-cm chamber on a 55°C hot plate (Bioseb) for 30 s. Each animal’s total number of nocifensive behaviors on the plate (paw flicks or licks, and jumps) was recorded.

### Pharmacological injections

Clozapine N-oxide (RTI), nitroglycerin (American Regent Inc.), cisplatin (West Ward) and saline or vehicle controls were injected at 10 mL/kg body weight. CNO was administered at 1 mg/kg for hM3Dq experiments and 5 mg/kg for hM4Di experiments, nitroglycerin at 10 mg/kg, cisplatin at 2.3 mg/kg and saline at 0.9% sodium chloride; vehicle for nitroglycerine was 6 % propylene glycol 6 % ethanol and saline. Lithium chloride (Fisher Scientific) and the saline control for this condition were injected at 15 mL/kg. Lithium chloride was administered at 0.2 M and the saline was 0.9% sodium chloride. All paw-withdrawal measurements were made 2 h after each CNO injection and daily thereafter unless otherwise indicated.

### Optogenetic stimulation

After recovery from surgery, mice were acclimated to fiber-optic cable attachment. For von Frey, allodynia assessment, light-pulse trains (20 Hz; 2 s “on”, 2 s “off”) were delivered for 5 or 20 min as described in the text. Stimulation paradigms were programmed using a Master8 (AMPI) pulse stimulator that controlled a blue-light laser (473 nm; LaserGlow). The power of light exiting each side of the branching fiber-optic cable was adjusted to 10 ± 1 mW. All paw-withdrawal measurements were made 2 h after each ChR2 stimulation and daily thereafter

### Immunohistochemistry

Mice were anesthetized with phenobarbitol (0.2 ml, i.p.; Akorn) and perfused transcardially with phosphate-buffered saline (PBS) followed by 4% paraformaldehyde (PFA, Electron Microscopy Sciences) in PBS. Brains were post-fixed overnight in 4% PFA at 4°C, cryoprotected in 30% sucrose, frozen in OCT compound (ThermoFisher) and stored at -80°C. Coronal sections (30 µm) were cut on a cryostat (Leica Microsystems) and collected in cold PBS. For immunohistochemistry experiments, sections were washed three times in PBS with 0.2% Triton X-100 (PBST) for 5 min and incubated in blocking solution (3% normal donkey serum in PBST) for 1 h at room temperature. Sections were incubated overnight at 4°C in PBS with primary antibodies including: chicken-anti-GFP (1:10000, Abcam, ab 13970) or rabbit-anti-dsRed (1:1000, Tacara, ab 632496). After 3 washes in PBS, sections were incubated for 1 h in PBS with secondary antibodies: Alexa Fluor 488 donkey anti-chicken or Alexa Fluor 594 donkey anti-rabbit (1:500, Jackson ImmunoResearch). Tissue was washed 3 times in PBS, mounted onto glass slides, and coverslipped with Fluoromount-G (Southern Biotech). Fluorescent images were acquired using a Keyence BZ-X700 microscope. Images were minimally processed using ImageJ software (NIH) to enhance brightness and contrast for optimal representation of the data. All digital images were processed in the same way between experimental conditions to avoid artificial manipulation between different datasets.

### RNAscope *in situ* hybridization

Mice were anesthetized with phenobarbital (0.2 ml, i.p.) then decapitated. Brains were rapidly frozen on crushed Dry Ice. Coronal sections (20 μm) were cut on a cryostat (Leica Microsystems), mounted onto glass slides, and stored at –80°C. RNAscope fluorescent multiplex assay was performed following the manufacturer’s protocols (ACD Biotechne). Samples were taken from 2 males and 2 females in each group (3 and 30 day) and several levels of the PBN were imaged for each animal using a Keyence BZ-X710 microscope. Using the superior cerebellar peduncle (scp) as the center point, images were acquired at 20x in a 3×3 grid then stacked and stitched together using Fiji for an initial total of 116 PBNs. Images of probe staining within the four-channel sets were subtracted from one another using Fiji’s image calculator function to remove background autofluorescence and minimally processed to enhance brightness and contrast for optimal representation of the data. After image optimization, due to difficulty in getting precisely matching bregma levels during sectioning, PBN anatomy was evaluated using fiber-tract location and general structure to categorize the images into two groups. “Rostral” sections were defined by presence of the caudal part of the nucleus of the lateral lemniscus (NLL) and a more triangular shape of the lateral PBN (Bregma -4.95 to -5.15), and “middle” sections were defined by presence of the longer central part of the ventral spinocerebellar tract (sctv) and narrower oval appearance of lateral PBN (Bregma 5.15 to -5.35). Sections that were deemed to be further rostral and caudal of these two categories, and sections that had tissue damage in the PBN were removed for a final count of 89 PBN images. Images were imported into QuPath and a region of interest was drawn over the lateral PBN using the surrounding fiber tracts and brain structure as a guide. RNA expression was quantified by thresholding using the subcellular detection function in QuPath. Cells were deemed positive for *Calca* or *Cck* if they had 5 or more puncta and cells were deemed positive for *Fos* if they had 4 or more puncta because *Calca* and *Cck* had denser transcript labeling. An average of ∼1700 DAPI-positive cells were analyzed for each section for a total of ∼150,000 cells analyzed.

### Electrophysiology

Mice were deeply anesthetized with Euthasol (i.p. 1 µl per 10 g body weight) and intracardially perfused with ice-cold cutting solution containing (in mM): 92 N-methyl-D-glucamine, 25 D-glucose, 2.5 KCl, 10 MgSO_4_, 1.25 NaH_2_PO_4_, 30 NaHCO_3_, 0.5 CaCl_2_, 20 HEPES, 2 thiourea, 5 Na-ascorbate, 3 Na-pyruvate. Brains were quickly removed after perfusion and 250-μm coronal slices were prepared (Leica VT1200) in the same ice-cold solution. Brain slices were kept in the cutting solution at 33°C for 10 min and then transferred to a room temperature recovery solution containing (in mM): 13 D-glucose, 124 NaCl, 2.5 KCl, 2 MgSO_4_, 1.25 NaH_2_PO_4_, 24 NaHCO_3_, 2 CaCl_2_, 5 HEPES for at least 1 h. Slices were individually transferred to 33°C artificial cerebral spinal fluid containing (in mM): 11 D-glucose, 126 NaCl, 2.5 KCl, 1.2 NaH_2_PO_4_, 26 NaHCO_3_, 2.4 CaCl_2_, 1.2 MgCl_2_ for recording. All solutions were saturated with 95% O2/5% CO2 and adjusted to pH 7.3–7.4, 300–310 mOsm.

Epifluorescence microscope (OLYMPUS BX51WI) was used to visualize *Calca* neurons expressing AAV-DIO-hM3Dq-mCherry and AAV-DIO-mCherry. A 3-5 MΩ glass pipet containing (in mM): 135 K-gluconate, 4 KCl, 10 HEPES, 4 Mg-ATP, 0.3 Na-GTP (pH 7.35, 280 -300 mOsm) was used to record neuron activity. To record intrinsic firing frequencies, 800-ms current injections with 20-pA steps from -100 pA to 240 pA were applied in current clamp, with initial holding potential at -70 mV and repeated every 10 s. To record the efficiency of CNO application, cell-attached measurement was used to record action potential in voltage clamp with 0 pA holding current. CNO (3 µM) was bath applied after 3 min of action potential firing. All data were obtained using MultiClamp 700B amplifier (Molecular Devices). Data acquisition and analysis were done using pClamp 11 and Clampfit 11.0.3.

### Calcium imaging

AAV-Ef1a-DIO-GCaMP6m was injected into the PBN of *Calca^Cre/+^* mice. After 6 weeks of recovery, the nVista (Inscopix) microscope was attached and connected once a week to check the field of view. Recording was conducted using the IDAS program (Inscopix), and the best focus was determined through visual inspection. Occasionally, the mice received a brief air puff as an aversive stimulus. For multi-day imaging of *Calca* neurons, nVista was connected Ethovision (Noldus) via BNC cable to synchronize video recording and Ca^2+^ imaging to ensure that the onset and offset of the Ca^2+^ imaging session matched the video recording. The microendoscope was connected to a commutator (Inscopix) and attached to the baseplate on the mouse, which was then housed in an open-top cage for 4 days to maintain the same field of view throughout the experimental period.

On day 1, a vehicle solution (6 % propylene glycol 6 % ethanol and saline) was injected intraperitoneally, and the mouse was placed in the von Frey-stimulation chamber for a 45-min acclimation. The imaging session consisted of 5 min of basal activity and 10 min of von Frey stimulation. During the von Frey-stimulation sessions, the mouse was exposed to 8 stimulations of 0.4-g von Frey filament with 1-min, inter-trial intervals. After the imaging session, the LED was turned off, but the microendoscope remained connected, and the mouse was then returned to open-top cage until the start of the next day’s experiment. On day 2, NTG (10 mg/kg, dissolved in 6 % propylene glycol 6 % ethanol and saline) was injected intraperitoneally. The same imaging session as on day 1 was repeated. On days 3 and 4, the mice were placed in the von Frey-stimulation chamber for imaging as on previous days.

The imaging parameters had insignificant photobleaching, but sufficient fluorescence (LED power, 0.2 - 0.4; sampling rate 10 Hz). The raw data were processed using IDPS software (ver. 1.9.1, Inscopix). All images acquired over 4 days were concatenated for further analysis. After applying 4X spatial and 2X temporal downsampling, the data underwent spatial bandpass filtering to reduce background noise. Subsequently, motion correction was applied to the images based on a reference frame and region of interest (ROI) within the field of view (FOV). Using IDPS, ΔF/F movies were generated, then PCA/ICA analysis was performed to extract neuronal activities. In cases where there was significant motion or higher background noise, a manual ROI analysis method was employed. To do that, the maximum projection images were used as a reference for spatial information of the neurons, and ROIs were manually drawn based on the borders of the neurons. All neurons analyzed using PCA/ICA and manual ROI methods were visually inspected for each cell, taking into consideration their shape and dynamics, to ensure accuracy.

The outputs of PCA/ICA (ΔF/F) were processed using customized MATLAB code to calculate the Z-score. In the case of manual ROI analysis, ΔF/F was calculated as ΔF/F=(F−Fmean)/Fmean. Fmean indicates mean fluorescent during day 1 baseline. For basal activity analysis, concatenated Ca^2+^ traces were used to calculate Z-score, and the Z-scored data were compared across days using the formula: Z=(F−Fmean)/Fstd. Fstd indicates standard deviation of fluorescent during day 1 baseline. For the analysis of von Frey filament-elicited responses, the traces from -20 s to 20 s were extracted and used to calculate the Z-scores. To compare the fluorescent responses, all data points were calculated relative to the -20 s to 0 s periods because the mice exhibited different basal activities every day due to intraperitoneal injections or allodynia.

Responsive neurons were identified through statistical analysis comparing the area under the curve (AUC) of pre- and post-stimulation. The AUCs were calculated for three time blocks (-5 to 0 s, 0 to 5 s, and 5 to 10 s) across 8 trials and compared using the Wilcoxon Signed-rank test. Neurons were classified as “increased” if there was a significant increase in the AUC in more than one post-stimulation block. If neither post-stimulation block showed statistical significance, neurons were classified as “unresponsive”. Neurons that exhibited a statistically significant decrease in activity across two blocks were classified as “decreased”.

## QUANTIFICATION AND STATISTICAL ANALYSIS

Data were analyzed in GraphPad Prism 9.5.1 (Graphpad Software) by two-way ANOVA with Šidák post hoc correction; for longitudinal assays Bonferroni’s multiple comparisons correction was applied; *p* < 0.05 was deemed statistically significant. All data are presented as the mean ± standard error of the mean (SEM). The asterisks in the figures represent the p values of post hoc tests corresponding to the following values * p < 0.05, ** p < 0.01, *** p < 0.001, **** p < 0.0001. Following histology and imaging, any mouse whose targeted injection site was wrong was excluded from experimental analysis.

## SUPPLEMENTAL INFORMATION

Supplemental information can be found online at ____.

## ACKNOWLEDGMENTS

We thank Susan Phelps for maintaining the mice used in this study and our colleagues for their input during this study and preparation of the manuscript. Logan Condon was funded by NIH NIDA 1F30DA057845-01 and Tyler Nelson was funded by NIH NINDS K00NS124190. The Palmiter lab is funded by the Howard Hughes Medical Institute.

## AUTHOR CONTRIBUTIONS

L.F.C. and R.D.P. designed the study. L.F.C., S.Y., S.P., F.C. and J.L.P. performed experiments. T.S.N. assisted with experiments. L.F.C., S.Y, S.P., F.C. and J.L.P. analyzed data. L.F.C and R.D.P wrote the manuscript with input from all the authors.

## DECLARARTION OF INTERSTS

Authors declare no conflicting interests.

## INCLUSIVITY AND DIVERSITY

We support inclusive, diverse, and equitable conduct of research.

---

**Figure S1.**
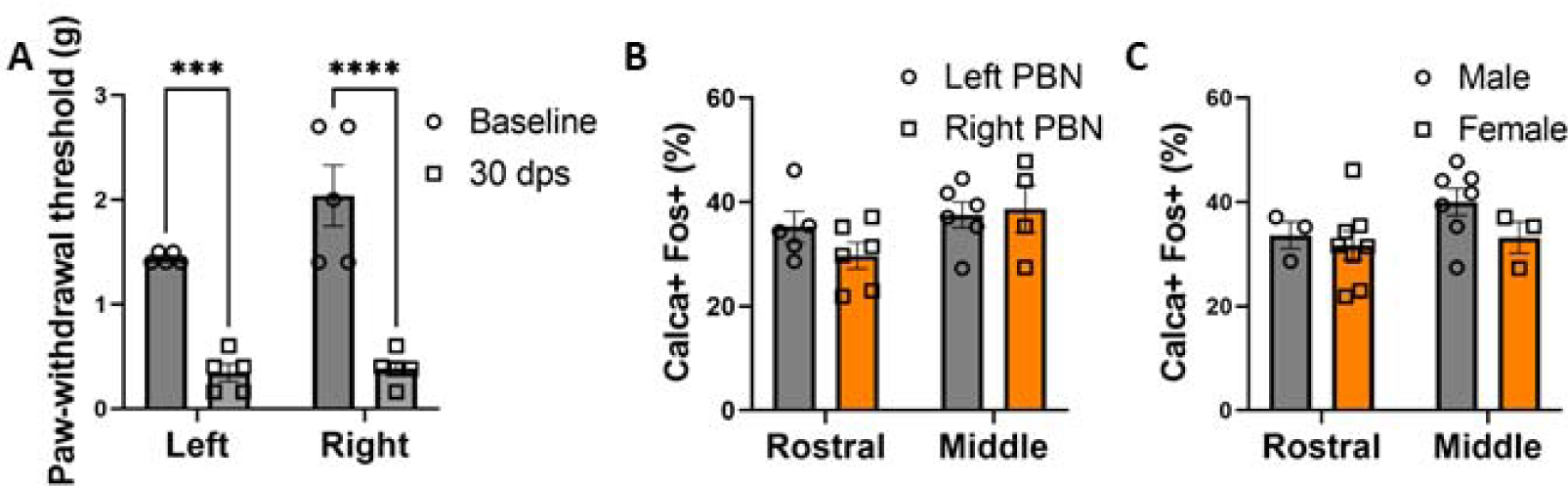
PBN *Calca* neurons bilaterally exhibit uniform activity and do not have sexual dimorphic activity. (A) pSNL resulted in bilateral allodynia, n = 5. (B) There was no difference between left and right PBN *Calca*-*Fos* colocalization 3 days post unilateral (left) pSNL. Rostral left PBN, n = 5; rostral right PBN, n = 6; middle left PBN, n = 6; middle right PBN, n = 4. (C) There was no difference between male and female PBN *Calca*-*Fos* colocalization 3 days post pSNL. Rostral male, n = 3; rostral female, n = 7; middle male, n = 7; middle female, n = 3.

**Figure S2.**
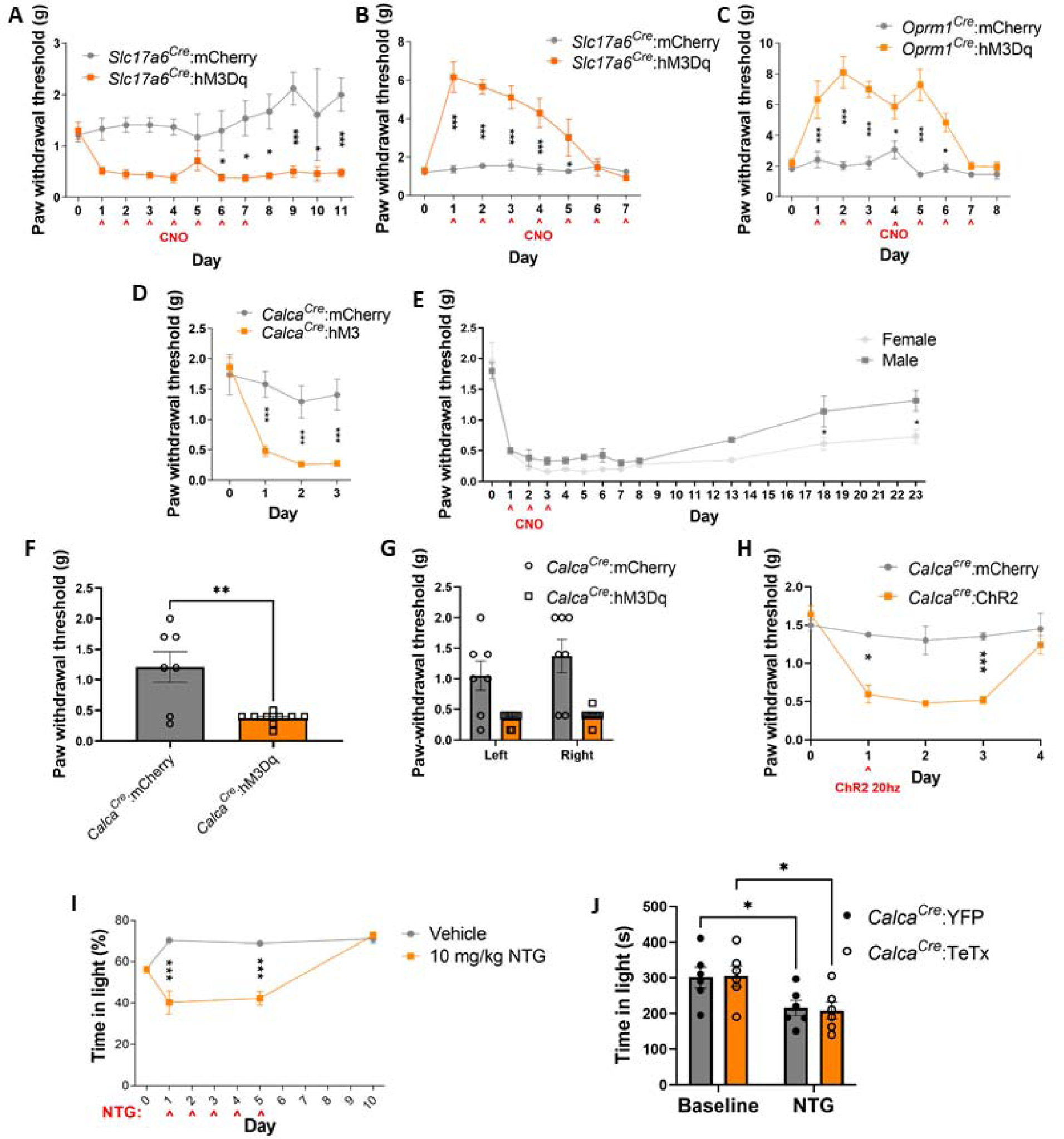
Additional behavioral findings. (A) Chronic stimulation of PBN *Slc17a6* (Vglut 2) neurons via hM3Dq and CNO resulted in persistent allodynia (measured 23 h post injection). (B) Stimulation of PBN *Slc17a6* neurons via hM3Dq and CNO resulted in analgesia 2 h post injection. (A-B) *Slc17a6^Cre^*:mCherry n = 5, *Slc17a6^Cre^*:hM3Dq n = 6. (C) Stimulation of PBN *Oprm1* neurons via hM3Dq and CNO resulted in analgesia 2 h post injection. *Oprm1^Cre^*:mCherry n = 5, *Oprm1^Cre^*:hM3Dq n = 7. (D) Stimulation of PBN *Calca* neurons via hM3Dq and CNO resulted in allodynia 23 h after CNO treatment. *Calca^cre/+^*:mCherry n = 5 and *Calca^cre/+^*:hM3Dq n = 7. (E) Stimulation of PBN *Calca* neurons via hM3Dq and CNO resulted in persistent allodynia in both male and female animals. Female, n = 3; male, n = 4. (F) Unilateral stimulation of PBN Calca neurons via hM3Dq and CNO resulted in allodynia. *Calca^cre/+^*:mCherry n = 7 and *Calca^cre/+^*:hM3Dq n = 10. (G) Unilateral stimulation of PBN Calca neurons via hM3Dq and CNO affected left and right hind paw withdrawal threshold equivalently. *Calca^cre/+^*:mCherry n = 7 and *Calca^cre/+^*:hM3Dq n = 10. (H) Unilateral stimulation of PBN *Calca* neurons via ChR2 and 473-nm light (20 min, 20 Hz, 2 s on 2 s off) resulted in persistent allodynia. *Calca^cre/+^*:mCherry n = 4 and *Calca^cre/+^*:ChR2 n = 4. (I) NTG injection resulted in photophobia that did not persist past the point of NTG administration. Vehicle, n = 4; NTG, n = 4. (K) TeTx expression in PBN *Calca* neurons did not prevent the development of NTG-driven photophobia. *Calca^Cre/+^*:YFP n = 6 and *Calca^Cre/+^*:TeTx n = 6.

**Figure S3.**
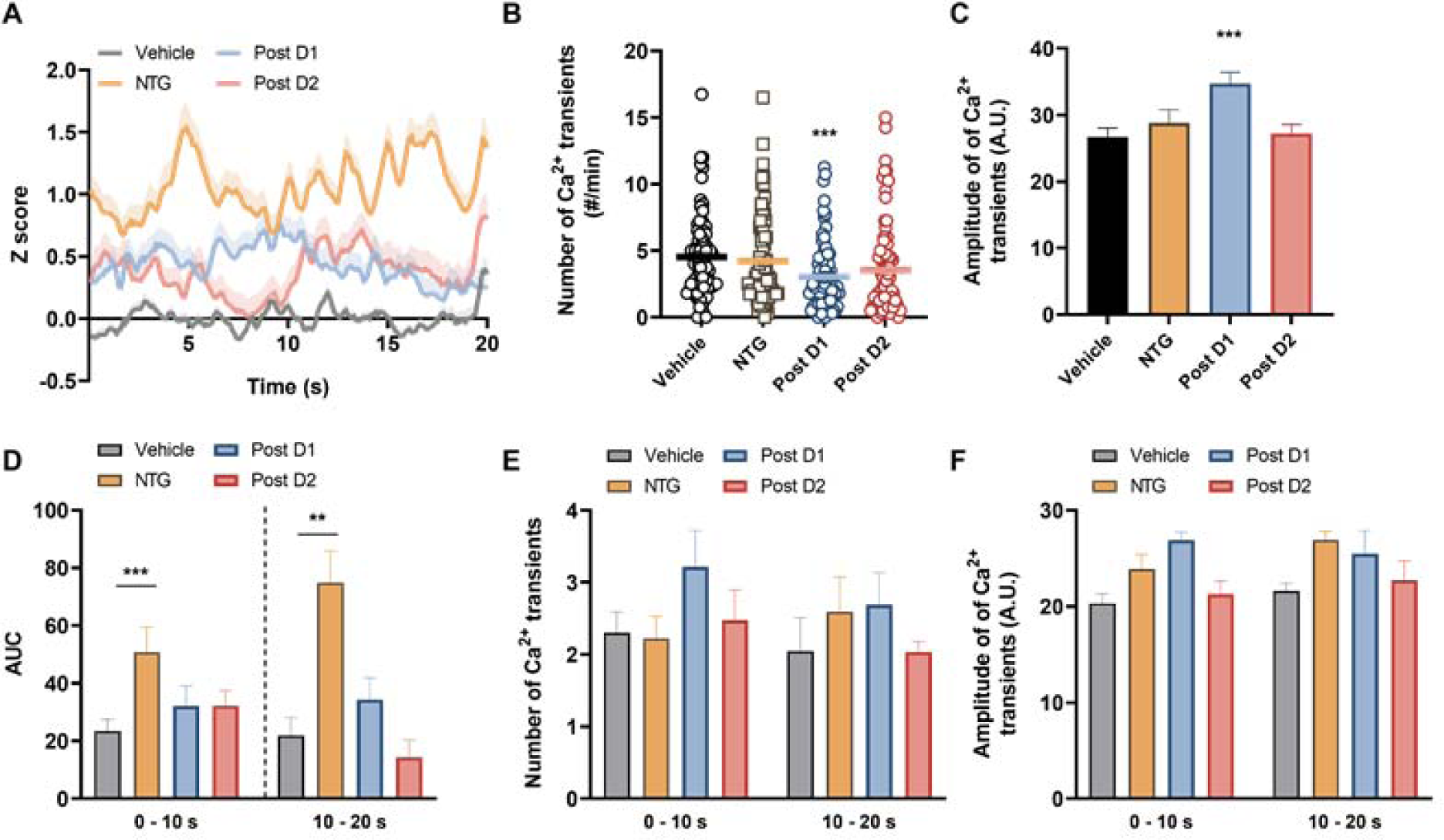
NTG injection does not affect the total number of or amplitude of calcium transients in *Calca* neurons. (A) Average traces of individual neurons during baseline period (10 min before von Frey filament application). Shaded area indicates. +S.E.M. (B) Number of Ca2+ transients during 10 min of baseline. (C) Amplitude of Ca2+ transients during 10 min of baseline. (D) Average AUC of individual neurons evoked by von Frey filament. Bar indicates mean S.E.M. (E) Number of Ca2+ transients during 10 min of baseline. (F) Peak amplitude during 20 sec of post stimulation period. (A-F) n = 3 animals, 79 neurons.

## Notes

### Competing Interest Statement

The authors have declared no competing interest.

